# Estimation of Kinematics from Inertial Measurement Units Using a Combined Deep Learning and Optimization Framework

**DOI:** 10.1101/2020.12.29.424030

**Authors:** Eric Rapp, Soyong Shin, Wolf Thomsen, Reed Ferber, Eni Halilaj

## Abstract

*The difficulty of estimating joint kinematics remains a critical barrier toward widespread use of inertial measurement units in biomechanics. Traditional sensor-fusion filters are largely reliant on magnetometer readings, which may be disturbed in uncontrolled environments. Careful sensor-to-segment alignment and calibration strategies are also necessary, which may burden users and lead to further error in uncontrolled settings. We introduce a new framework that combines deep learning and top-down optimization to accurately predict lower extremity joint angles directly from inertial data, without relying on magnetometer readings. We trained deep neural networks on a large set of synthetic inertial data derived from a clinical marker-based motion-tracking database of hundreds of subjects. We used data augmentation techniques and an automated calibration approach to reduce error due to variability in sensor placement and limb alignment. On left-out subjects, lower extremity kinematics could be predicted with a mean (*± *STD) root mean squared error of less than 1*.*27 ° (*± *0*.*38* °*) in flexion/extension, less than 2*.*52* ° (± *0*.*98 °) in ad/abduction, and less than 3*.*34* ° (± *1*.*02* °*) internal/external rotation, across walking and running trials. Errors decreased exponentially with the amount of training data, confirming the need for large datasets when training deep neural networks. While this framework remains to be validated with true inertial measurement unit (IMU) data, the results presented here are a promising advance toward convenient estimation of gait kinematics in natural environments. Progress in this direction could enable large-scale studies and offer an unprecedented view into disease progression, patient recovery, and sports biomechanics*.

## 1. Introduction

The ability to passively estimate movement kinematics in natural environments with inertial measurement units (IMUs) could transform how we monitor, diagnose, and treat mobility limitations. These devices are now unobtrusive, can flex and bend with the skin, and allow for patient monitoring throughout the day (Patel *et al*., 2012 [18]; Shull *et al*., 2014 [31]; Shull and Damian, 2015 [30]; Son *et al*., 2014 [32]). Despite recent progress in hardware miniaturization, turning large multimodal data from wearable sensors into meaningful biomechanical outcomes that can be easily interpreted in the context of healthy and pathological movement remains a key challenge toward their widespread use (Picerno, 2017 [20]). Biomechanists and rehabilitation specialists have traditionally characterized gait using joint kinematics and kinetics. While segment inertial data generated by IMUs may also lead to important actionable insights, accurate estimation of joint angles is needed to place future findings in the context of past work.

There are currently no streamlined tools to accurately estimate three-dimensional (3D) joint kinematics from wearable sensors worn in uncontrolled environments. Strapdown integration of inertial data introduces drift, and sensor fusion algorithms that rely on magnetometer data suffer from ferromagnetic disturbances (de Vries *et al*., 2009 [3]). Solutions that incorporate full-body biomechanical models (Robert-Lachaine *et al*., 2017a [24], 2017b [25], 2020 [26]) are currently not portable for anytime, anywhere use, and accuracy over long durations remains to be demonstrated. Additionally, the dependence of most algorithms on accurate sensor-to-segment alignment makes translation difficult for multi-day monitoring outside of the laboratory, given human error in sensor placement. Static and dynamic calibrations (Cutti *et al*., 2010 [2]; Favre *et al*., 2009 [5]; Picerno *et al*., 2008 [21]; Roetenberg *et al*., 2009 [27]) may also add to lack of compliance and increased drop-out rates in remote monitoring studies. Further, static poses or functional calibration trials may not be performed as expected in remote scenarios, especially when users suffer from a mobility-limiting condition.

Deep learning and iterative optimization techniques offer a new opportunity to overcome the limitations of previously proposed approaches for estimating kinematics from IMUs. Deep neural networks are highly efficient in learning non-linear relationships from high-dimensional data, such as dense time series from wearable sensors. A key drawback, however, is that they require large datasets to generate accurate models, and such datasets are scarce in the field of biomechanics. A large dataset that contains both IMU data and ground truth joint kinematics from marker-based motion tracking systems, for example, is currently not available to the research community. In other domains, however, synthetic data have been successfully used to train accurate predictive models (Jaderberg *et al*., 2014 [10]). Additionally, optimization approaches have demonstrated success in improving pose estimation in computer vision applications (von Marcard *et al*., 2018 [34]).

The goal of this study was to build deep learning models for predicting 3D lower extremity joint kinematics directly from angular velocity and linear acceleration data, circumventing the limitations of magnetometer-dependent algorithms. To generate sufficient data to train such models, we created synthetic inertial data from a marker-based motion capture database of hundreds of subjects collected at a clinical center (Ferber *et al*., 2014 [6]; Osis *et al*., 2015 [17]). We further incorporated data augmentation strategies (Shorten and Khoshgoftaar, 2019 [29]) to increase the effective sensor placement variability represented in the data, allowing the developed models to tolerate placement ambiguity without compromising accuracy. Additionally, we used an iterative top-down optimization approach to improve the predictions of the deep learning models.

## 2. Methods

### 2.1. Data Collection and Pre-processing

To train the models, we used marker-based motion capture data that were previously collected at the University of Calgary Running Injury Clinic after receiving approval from the University of Calgary’s Conjoint Health Research Ethics Board (Ferber *et al*., 2016 [7]; Jauhiainen *et al*., 2020 [11]; Phinyomark *et al*., 2018 [19]; Pohl *et al*., 2010 [22]). Retro-reflective marker trajectories were collected at 200 Hz using eight high-speed infrared video cameras (Vicon Motion Systems Ltd., Oxford, UK). After a static neutral trial, subjects performed 60-second walking and running trials at self-selected speeds on a treadmill, following a 2–5 minute acclimation period. The participants were patients who enrolled in either clinical or research activities and gave written informed consent prior to participation. A number of the participants were pain-free, while others were experiencing a lower extremity running-related injury.

To perform 3D kinematic analysis, segment coordinate systems were constructed based on anatomical markers placed on the following landmarks: 1st and 5th metatarsal heads; medial and lateral malleoli; medial and lateral femoral condyles; greater trochanter (bilateral); anterior superior iliac spine (bilateral); iliac crest (bilateral). Tracking marker clusters were placed on the pelvis and bilateral thighs and shanks. A rigid shell with three markers was placed over the sacrum with the two superior markers at the level of the posterior superior iliac spines, while rigid shells with four markers were attached to the shank and thigh (Fig. 1A). Tracking markers for the feet were placed on the posterior aspect of the shoes: two markers were vertically aligned on the posterior heel counter with a third marker placed laterally. Marker trajectories were filtered with a 10 Hz low-pass 2nd order recursive Butterworth filter. Joint angles were calculated with custom software (Running Injury Clinic Inc., Calgary, Alberta, Canada) using the six degreeof-freedom approach and expressed in terms of joint coordinate systems recommended by the International Society of Biomechanics (Grood and Suntay, 1983 [8]; Wu and Cavanagh, 1995 [35]). Walking trials from 420 subjects (203 male and 217 female, height of 172.53 ± 8.94 cm, weight of 71.24 ± 12.71 kg) and running trials from 580 subjects (292 male and 288 female, height of 172.70 ± 11.39 cm, weight of 71.30 ± 12.97 kg) were selected for further analysis here, after excluding data with marker-tracking errors.

**Figure 1.**
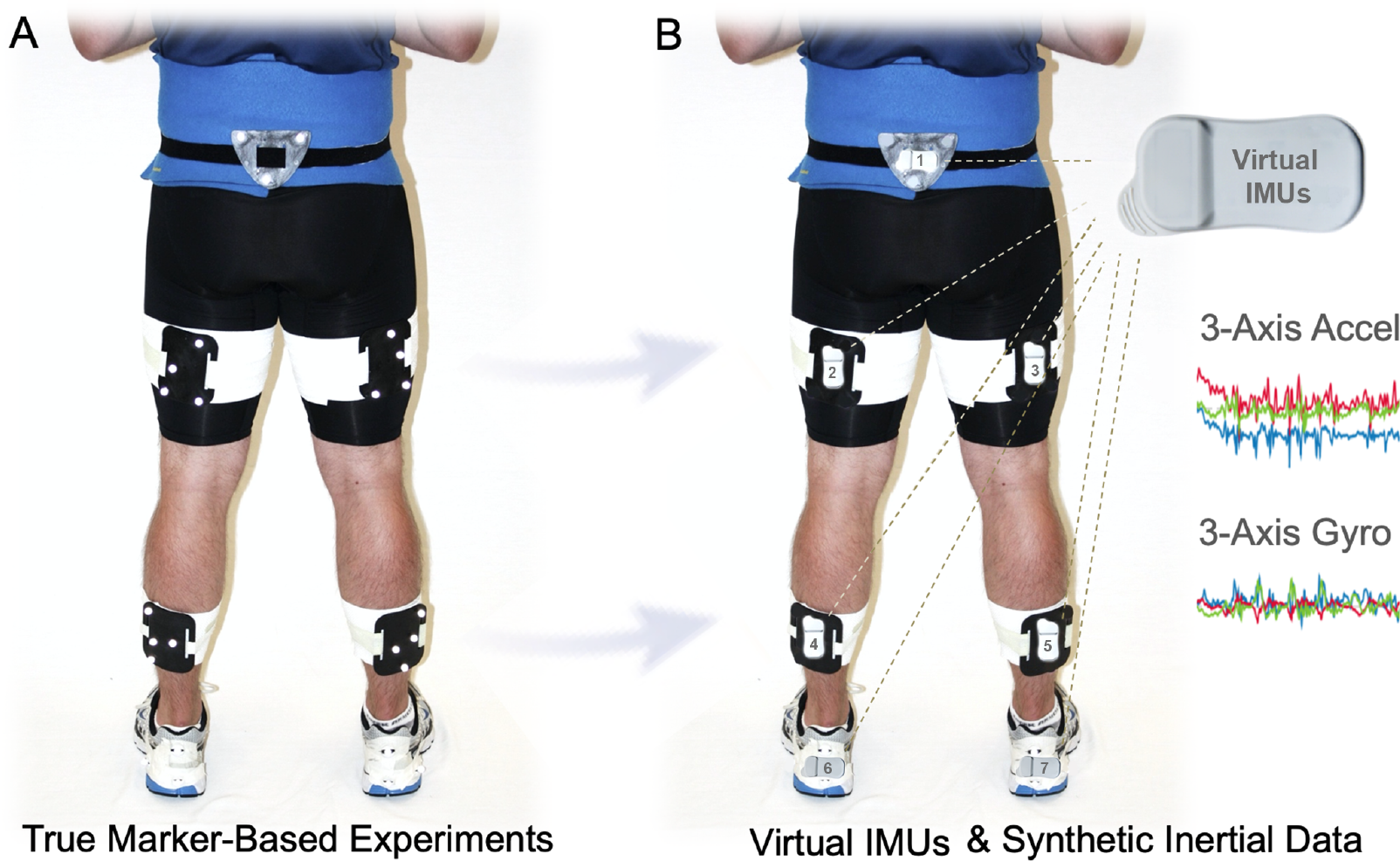
Experimental Set Up and Data Generation. A) Marker-based motion capture data previously collected at the Calgary Running Injury Clinic from hundreds of subjects. B) Tracking marker clusters were used to place virtual IMUs in the sacrum, thighs, shanks, and feet (1 − 7, respectively). Synthetic accelerometry and gyroscope data were generated by taking numerical derivatives and adding Gaussian noise.

We generated synthetic inertial data by placing virtual sensors on the tracking marker clusters (Fig. 1B). Seven coordinate systems were created for the lower body (pelvis and both thighs, shanks, and feet) based on the marker clusters, with the origins located at the centroids of the clusters. Inertial data—angular velocity and linear acceleration—were generated through numerical differentiation. Linear acceleration was transformed from a global to a local coordinate system after adding gravity, to match true accelerometer readings. Gaussian noise with a standard deviation that was 15

### 2.2. Deep Neural Networks for Segment Pose and Joint Angle Prediction

We built deep neural networks that used the synthetic inertial data described above (angular velocities and linear accelerations) from two adjacent segments to predict true kinematics for a particular joint (Fig. 2). The networks were initially trained to predict both segment orientation and joint angles. Each model took the simulated inertial data from the two adjacent segments as input and predicted both adjacent segment orientations and 3-D joint angles. We tested two overarching deep learning architectures with complementary strengths. The first was a one-dimensional convolutional neural network (Conv1D), which makes efficient predictions on a time-window that slides over the entire time series and therefore is more efficient for offline processing. The second was a long short-term memory network (LSTM), which is a specific recurrent neural network that makes predictions one frame at a time, updating an internal state to propagate previous information. LSTMs trade lower computational efficiency during training for higher efficiency in real-time applications.

**Figure 2.**
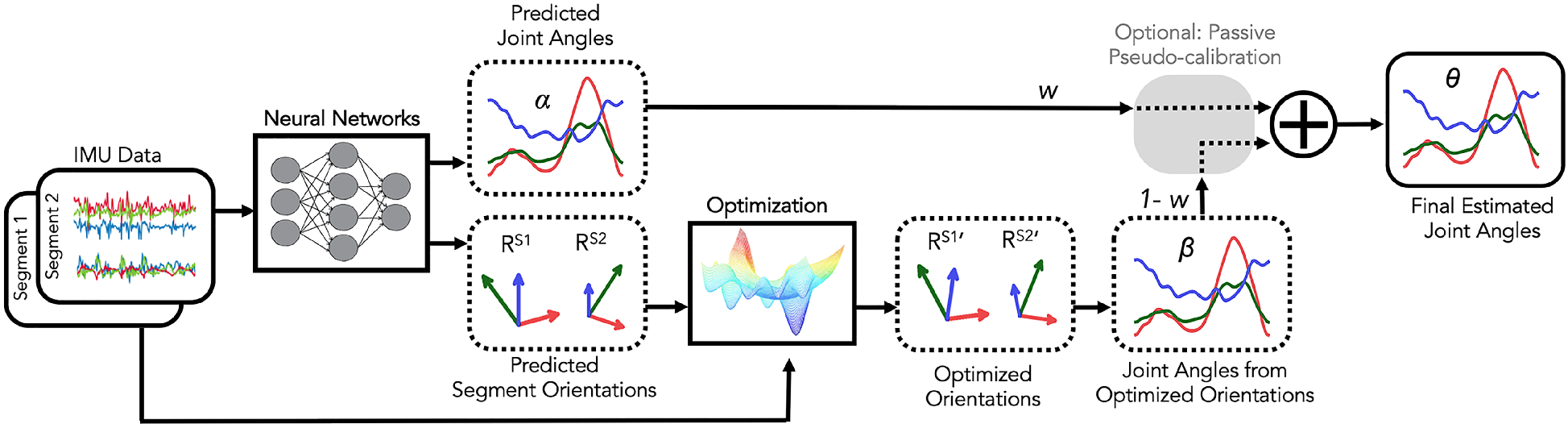
Hybrid Deep Learning and Optimization Approach. The synthetic sensor data (tri-axial linear acceleration and angular velocity) from adjacent segments were used to predict true segment orientation 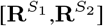 and joint angles [*α*_1_,*α*_2_,*α*_3_]. Segment orientation predictions were updated iteratively (optimization step) until the angular velocity associated with them best matched the virtual sensor data, and updated joint angles were computed from these optimized orientations [*β*_1_,*β*_2_, *β*_3_]. The final 3D joint angles (***θ***) were then computed using a weighted average of the prediction from the deep learning model (***α***) and the results of the optimization routine (***β***).

The learning rate, learning rate decay factor, number of batch iterations, dropout, and the number and size of the hidden layers for both the Conv1D and LSTM models, as well as window size for the Conv1D models and directionality for the LSTM models, were tuned using a Tree of Parzen Estimators optimization algorithm (Bergstra *et al*., 2011 [1]). For each set of hyperparameters, we trained the model by randomly selecting a sequence length of 1 second in stage one and a sequence length of 2 seconds in stage two. This two-stage scheme allowed the model to update its parameters by first learning the general trends of the gait cycle and then the cyclical nature of the whole time series.

### 2.3. Data Augmentation

To assess if data augmentation approaches could improve sensitivity to sensors placement variability, we performed additional experiments. We initially aligned the virtual sensors with the marker clusters and trained models using only these sensor placement configurations (baseline model). The data were then augmented to account for sensor orientation variability by drawing randomly sampled rotations from a normal distribution (STD = 13.5 °) centered around the “true” orientation via an online augmentation paradigm (Krizhevsky *et al*., 2012 [14]; Shorten and Khoshgoftaar, 2019 [29]). Models were also trained with the augmented dataset (augmented model). We tested the performance of both of these models on test data that were not augmented. Additional experiments were run to gauge the ability of each of these models to compensate for variability in sensor placement for the test subjects, which was achieved by augmenting the test data in a similar manner to the training data.

### 2.4. Segment Pose Optimization

To further improve accuracy, we implemented a top-down optimization method that fine-tuned the neural network prediction ***α*** by comparing the angular velocities (**Ω**^*′*^) from the predicted segment orientations **R**^**S**^ with the angular velocities from the true virtual IMU data (**Ω**). Each body-segment orientation was updated iteratively until the angular velocities associated with it best matched the virtual IMU data (Fig. 2). The final 3D joint angles (***θ***) were then computed using a weighted average of the prediction from the deep learning model (***α***) and the results of the optimization routine (***β***). Here, we refer to this as the optimized model.

Specifically, the combined deep learning and optimization approach includes two stages: initialization and frame-by-frame optimization. At the first stage, the deep neural networks estimate an initial joint angle (***α***) and adjacent body segment orientations 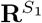 and 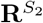. The frame-by-frame optimization stage updates the predicted segment orientations iteratively by minimizing the following error term for each frame, *k*:

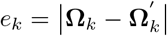

The optimization was carried out using an unconstrained Broyden-Fletcher-Goldfarb-Shanno optimizer with a function tolerance of 10^*-*6^, gradient tolerance of 10^*-*5^, and maximum iteration number of 200. To derive the final joint angle estimation ***θ***, we used a weighted average of the neural network prediction ***α*** and the optimization result ***β***, where ***θ* = *w · α* + (1 *-w*) *· β***. The optimal weight w was learned by minimizing the error between ***θ*** and the ground truth joint angles from optical motion tracking on the training dataset.

### 2.5. Passive Pseudo-calibration

Since the approach is purely data-driven and does not require a biomechanical model with new data, an active calibration routine with either static poses or dynamic functional tasks is not necessary. Data-driven models, however, generalize better to subjects whose neutral limb alignment is most represented in the training dataset. To overcome this limitation, we expressed the time series as a function of the mean joint angle in each degree of freedom. We call this procedure passive pseudo-calibration, since it does not engage the user and is not meant to replace a standard calibration. It does, however, ensure that subjects with limb alignments away from the mean have the same chance at accurate prediction of kinematics as subjects who are closer to the mean.

### 2.6. Performance Evaluation

The data were split into training (80%), validation (10%), and test (10%) sets, and separate predictive models were trained for each joint. The root-mean-squared error (RMSE) between the predicted and ground truth kinematics was computed for each joint and each hyper-parameter configuration. The model with the minimum RMSE on the validation data was selected as the best model for each joint. Final models were then tested on the left-out, test data. Here we report RMSEs for the test data, using the best models trained with non-augmented data (baseline model) and augmented data (augmented model) before the optimization step, and after incorporation of the optimization step (optimized model). These models were evaluated with and without the passive pseudo-calibration step. To determine the effect of training data size on model performance, we trained and evaluated models using only 10, 50, 100, 200 subjects, in addition to the standard experiments with 80% of the total subjects. We randomly selected the subsets and repeated the procedure 5 times. Here we report mean (± STD) RMSEs from the 5 trials using models trained with augmented data and tested with non-augmented data, without implementing the optimization or pseudo-calibration steps since the goal was to determine the effect of sample size on the neural networks. All the models were tested on the same set of test subjects (*n* = 42 for walking and *n* = 58 for running) per trial. Repeated measures analyses of variance (ANOVAs) were carried out to determine if the RMSE changed significantly with more training data.

## 3. Results

Walking kinematics could be predicted with a mean (± STD) RMSE of less than 2.75 ° (± 0.66 °), while running kinematics could be predicted with a mean RMSE of less than 3.34 ° (± 1.02 °) using the optimized model, along with pseudo-calibration. During walking, flexion/extension was the most accurate degree of freedom, with a mean RMSE of less than 0.97 ° (±0.38 °) across the ankle, knee, and hip joints, followed by ab/adduction with a mean RMSE of less than 2.16 ° (± 0.85 °) and internal/external rotation with a mean RMSE of less than 2.75 ° (± 0.66) (Table 1; Fig 3). Similarly, during running, flex-ion/extension was the most accurate degree of freedom, with a mean RMSE of less than 1.27 ° (± 0.40 °) across the ankle, knee, and hip joints, followed by ab/adduction with a mean RMSE of less than 2.54 ° (± 0.98 °), and internal/external rotation with a mean RMSE of less than 3.34 ° (± 1.01 °) (Table 2; Fig 3). Generally, Conv1D models with 2-3 hidden layers, more than 50 nodes per layer, a window size greater than 55 ms, learning rate greater than 0.001, and 5000 iterations performed the best.

**Table 1.**
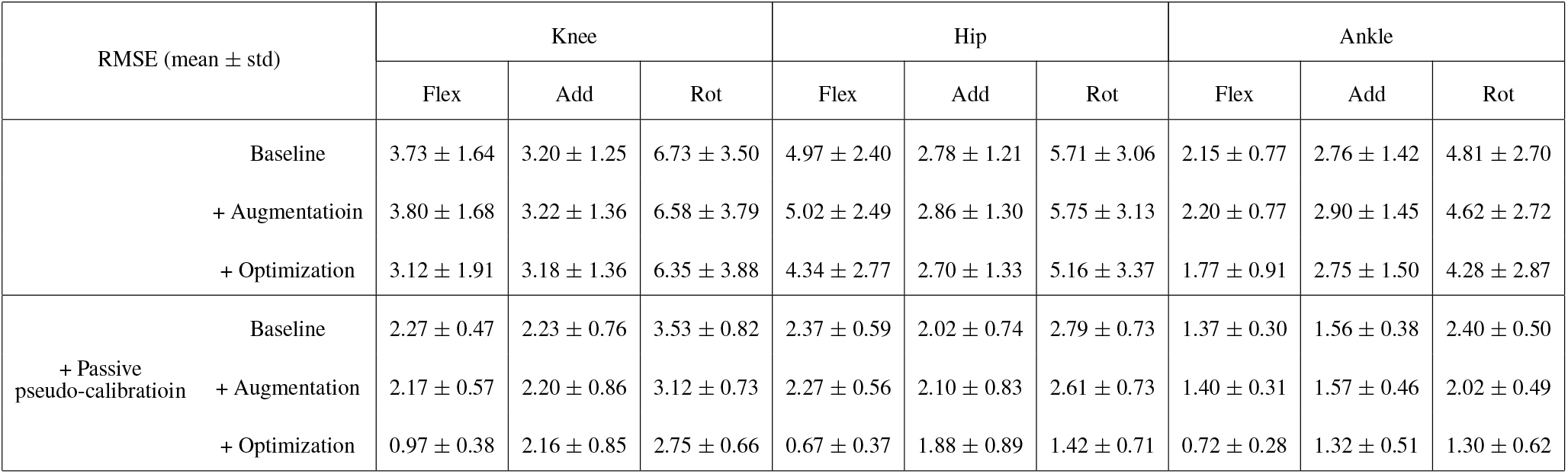
Predictive accuracy (mean RMSE ± STD in degrees) for walking kinematics at different steps of the pipeline.

**Table 2.**
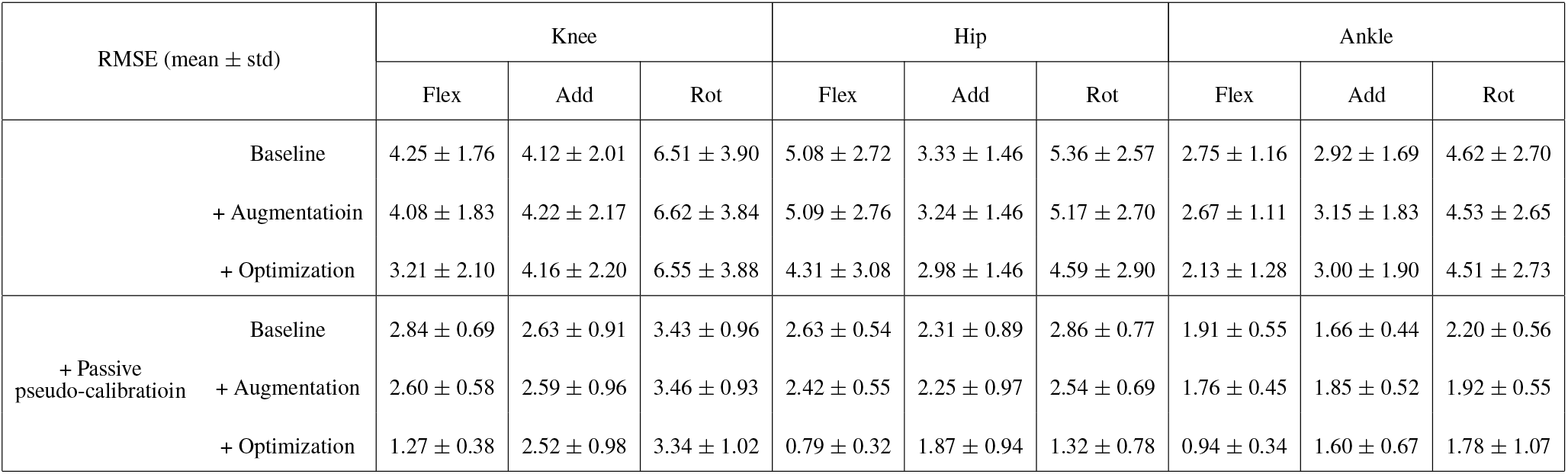
Predictive accuracy (mean RMSE ± STD in degrees) for running kinematics at different steps of the pipeline.

**Figure 3.**
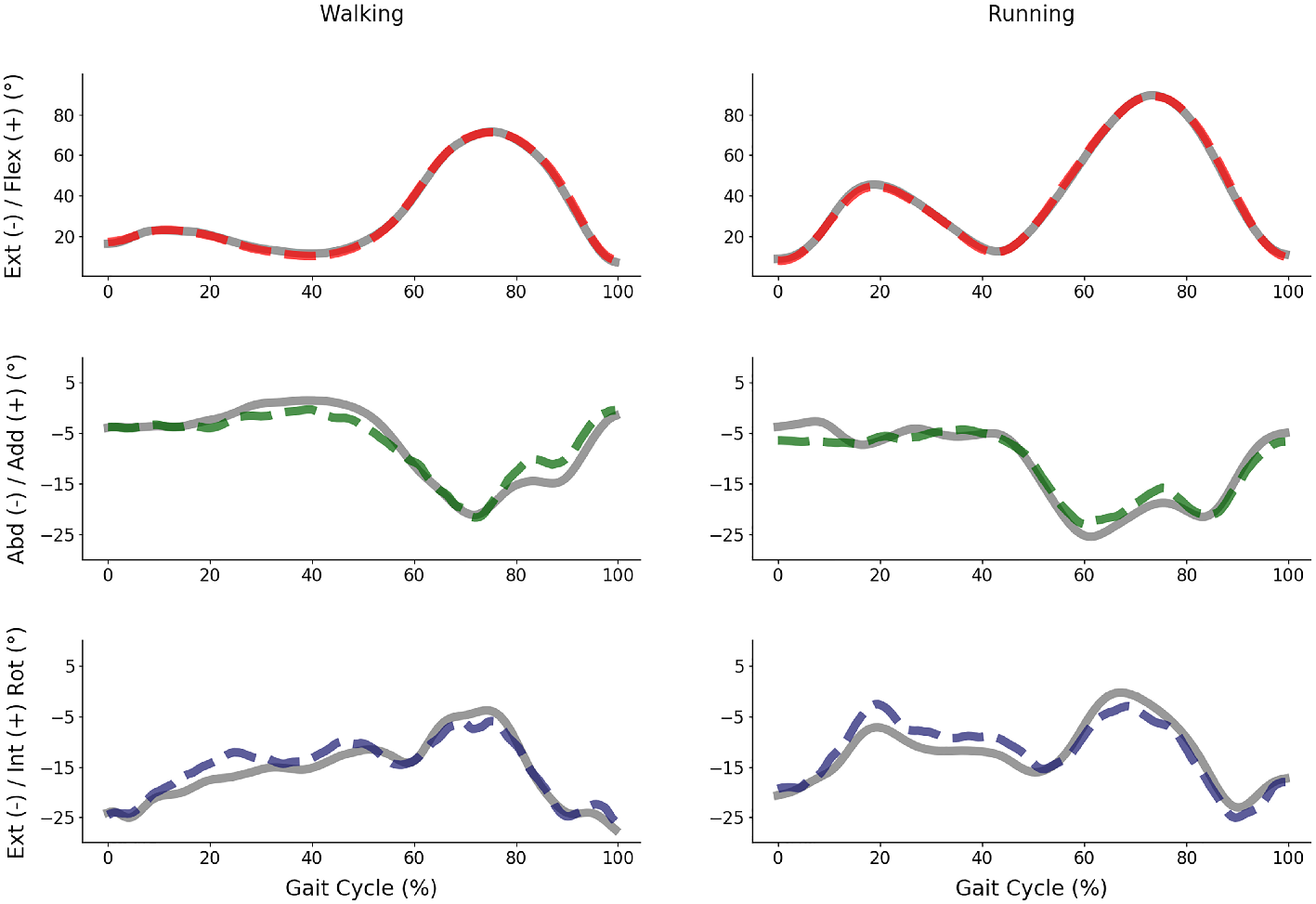
Predicted and Marker-Based Kinematics. A representative gait cycle from a subject whose root mean squared error of the predicted versus marker-based kinematics were close to the mean across all subjects. Predictions from the combined deep learning and optimization framework (dashed) closely match marker-based kinematics (solid gray), especially in flexion/extension (red), followed by add/abduction (green) and internal/external rotation (blue).

The final predictive performance reported above improved from the baseline neural network predictions through the use of iterative optimization, mainly for flexion/extension angles, but not the other degrees of freedom. Optimization improved prediction of knee flexion by over 15%, hip flexion by over 10%, and ankle flexion by over 15% (Tables 1 and 2). When passive pseudo-calibration was also applied, optimization had an even larger effect on final kinematics. In this case, prediction of knee flexion improved by over 50%, hip flexion by over 65%, ankle flexion by over 45%, and internal/external rotation of the hip and ankle by about 30% compared to the neural network outputs. For other degrees of freedom, optimal weight priors (***w***) were close to 1, indicating that the prediction was predominantly based on the neural network. Augmenting the training dataset to account for sensor application variability resulted in lower RMSEs when the test data were also augmented (Table 3), but not when the test data were not augmented (Table 3).

**Table 3.** Predictive accuracy (mean RMSE± STD in degrees) of the baseline and augmented models. No optimization or passive pseudo-calibration was applied.

| RMSE (mean ± std) | Knee | | | Hip | | | Ankle | | |
| --- | --- | --- | --- | --- | --- | --- | --- | --- | --- |
|  | Flex | Add | Rot | Flex | Add | Rot | Flex | Add | Rot |
| Walking Baseline | 8.27 ± 4.38 | 4.87 ± 2.01 | 9.38 ± 4.72 | 9.57 ± 5.70 | 4.33 ± 2.07 | 8.76 ± 4.92 | 5.79 ± 2.57 | 4.82 ± 2.34 | 7.12 ± 3.42 |
| + Augmentatioin | 3.87 ± 1.69 | 3.35 ± 1.43 | 6.77 ± 3.85 | 5.13 ± 2.41 | 2.82 ± 1.26 | 5.91 ± 3.25 | 2.32 ± 0.78 | 2.98 ± 1.48 | 4.70 ± 2.66 |
| Running Baseline | 7.56 ± 4.03 | 5.55 ± 2.72 | 8.54 ± 3.90 | 7.91 ± 4.29 | 4.45 ± 1.79 | 6.94 ± 3.15 | 6.64 ± 3.75 | 5.42 ± 2.50 | 7.09 ± 3.02 |
| + Augmentation | 4.23 ± 1.78 | 4.26 ± 2.18 | 6.59 ± 3.63 | 5.14 ± 2.72 | 3.37 ± 1.56 | 5.39 ± 2.69 | 2.82 ± 1.24 | 3.28 ± 1.88 | 4.60 ± 2.67 |

Increasing the amount of training data led to lower mean RMSEs when predicting kinematics in new subjects, during both walking and running (Fig. 4; *p* < 0.0001). Results from models using the maximum number of training subjects are different from those reported earlier (Table 1 and 2) since here we averaged across 5 trials. Also, the reported RMSEs do not include optimization and pseudo-calibration steps. Overall, fitted learning curves indicated improvements of over 1 degrees for flexion/extension. On a linear scale, the learning curves also started to, but did not completely, plateau as the training data increased from tens to hundreds of subjects, indicating that inclusion of more data in the future could continue to reduce the RMSE. Projections indicate that thousands of subjects are needed to improve the models by another degree (Fig. 4).

**Figure 4.**
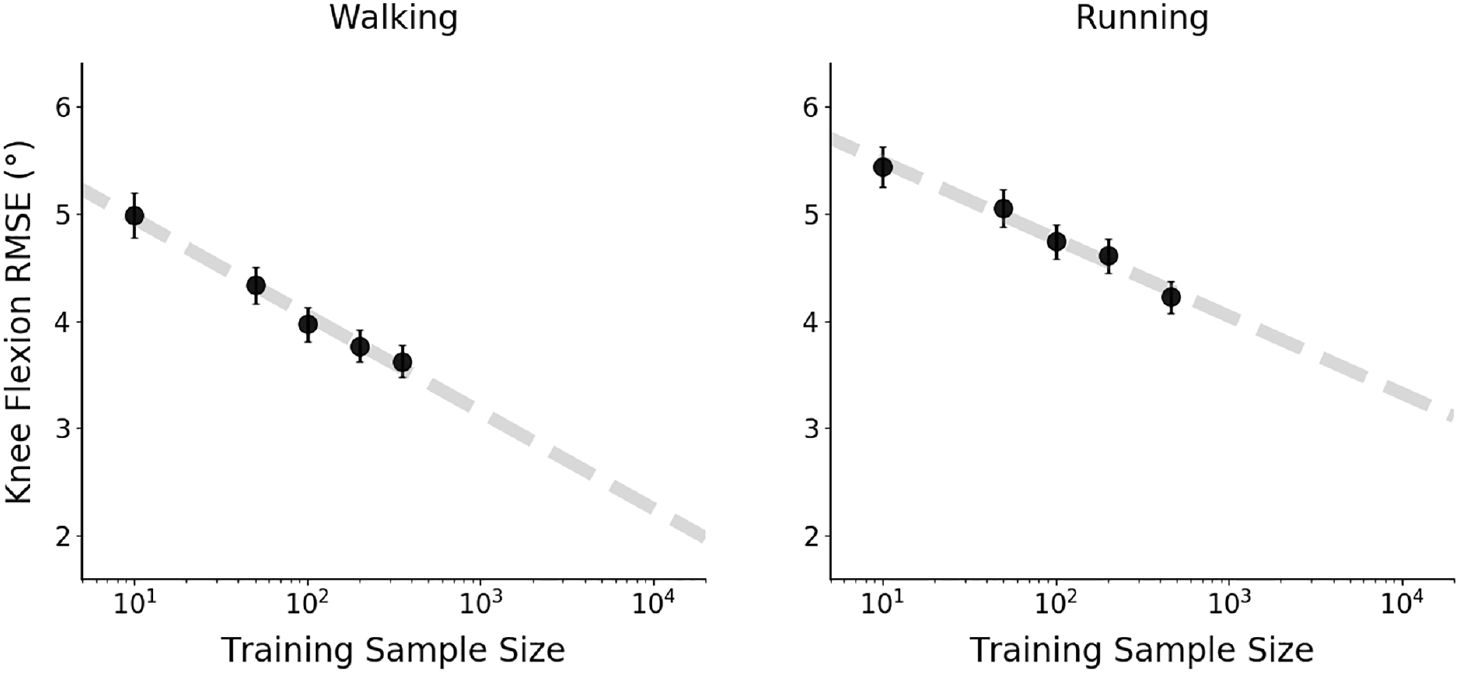
Neural Network Learning Curves. Models trained using a training data of different sizes (10, 50, 100, 200, and 80% of all the subjects) demonstrate that performance improves with larger datasets. Mean (± 95% CI) root mean squared error (RMSE) on the same set of left-out test data significantly decreased with models trained on larger data. Fitted lines (dashed gray) estimate the amount of training data needed to improve performance further.

## 4. Discussion

The goal of this study was to introduce a new framework that combines deep neural networks with top-down optimization for prediction of lower extremity kinematics from inertial sensing data. Using this hybrid approach and synthetic inertial data from a large clinical motion capture database, we could predict walking and running kinematics with accuracies that are similar to the reliability of marker-based motion tracking. We also demonstrated that augmentation techniques that increase effective sensor placement variability can improve model generalizability to sensor placement inconsistencies. Specifically, kinematics could be predicted with a mean RMSE of less than 1.27 (± 0.38 °) in flexion/extension, less than 2.52 ° (±0.98 ° ) in ad/abduction, and less than 3.34 ° (± 1.02 °) internal/external rotation. Predictive accuracy was consistent across male and female subjects and did not vary with subject weight and height (analysis not presented here). The higher RMSE in internal/external rotation may be explained by the lack of magnetometer data, which, when undisturbed, contribute toward heading estimation. Further, optical motion tracking, which was used as ground truth here, exhibits similar accuracies across degrees of freedom (Miranda *et al*., 2013 [16]; Tsai *et al*., 2011 [33]).

These promising results should be interpreted in the context of several limitations associated with the study. First, while we used data from hundreds of subjects, which contributed to improved model generalizability, the data were not acquired from true inertial sensors. Synthetic inertial data were generated from virtual sensors placed in accordance with optical motion tracking markers. We modeled the white noise associated with inertial sensors synthetically, but several other sources of noise, including zero-offset bias and soft tissue movement, were not modeled. While we expect the bias not to significantly affect the performance of these models, soft-tissue artifact remains a challenge. A second limitation is that, although moderate placement errors are surmountable given our data augmentation approach, the models developed here can only lead to plausible predictions if the sensors are placed on the same location as the location that we placed the virtual IMUs, which coincided with the location of the tracking marker clusters. Future work can focus on training more generalizable models that include virtual IMUs placed on several locations across each body segment. Another limitation is that the models were trained using treadmill walking and running data, while their intended use is for over-ground activities. Previous studies have documented little to no differences between treadmill and level ground walking and running (Riley *et al*., 2007 [23]). While we expect the models to generalize well to level walking and running, rugged terrain may introduce challenges. It is also important to note that for the proposed algorithms to be successful at predicting kinematics, walking and running should be segmented out of time series collected in free-living environments, which may include other activities and turns. Activity classification algorithms can predict walking and running with high accuracy given a few sensors worn on lower extremity segments (Mannini and Sabatini, 2010 [15]).

While validation with true sensor data is a necessary follow-up step, the accuracy of the presented framework is a promising advance toward prediction of joint kinematics from IMUs without relying on drift-prone algorithms and error-prone calibration techniques. Previously proposed approaches have reported RMSEs of more than 3 ° over limited durations (Dorschky *et al*., 2019 [4]; Karatsidis *et al*., 2018 [13]; Robert-Lachaine *et al*., 2020 [26], 2017b [25]). Because they rely on the use of magnetometers and filtering approaches, long-term reliability remains a limitation for natural environment applications. Algorithms that incorporate resetting to reduce drift (e.g., zero velocity update) may not generalize well to clinical populations with unpredictable gait patterns (Yang *et al*., 2013 [36]). In addition to not experiencing drift, our models are also well-suited for real-time applications that integrate monitoring with biofeedback. While the optimization step may introduce delays for real-time estimation and be better suited for post-hoc analyses, deep neural networks without the optimization step demonstrate mean RMSEs of less than 3.46 ° (± 0.93 °).

Reducing model sensitivity to sensor placement variability is desired for translation of wearable sensors out of the laboratory and clinic, where patients will have to apply them on their own, and likely more than once. While previous approaches are reliant on careful sensor-to-segment alignment and calibration trials, including functional (Favre *et al*., 2009 [5]), positional (Robert-Lachaine *et al*., 2017b [25]), and constraint-based (Seel *et al*., 2014 [28]) calibrations, our approach foregoes any active calibration and can overcome the challenges posed by sensor placement variability or need for supervision by trained research or clinical staff. We instead used data augmentation and passive calibration to overcome sensor placement and natural limb alignment variability. Data augmentation is an advantageous technique for increasing the amount of effective training data, and here we demonstrated that it can reduce the effect of sensor placement variability. Further, the predictive accuracy of our algorithms can be tuned passively after a few gait cycles (Fig. 5), without requiring attention from the user as is the case with active calibration routines.

**Figure 5.**
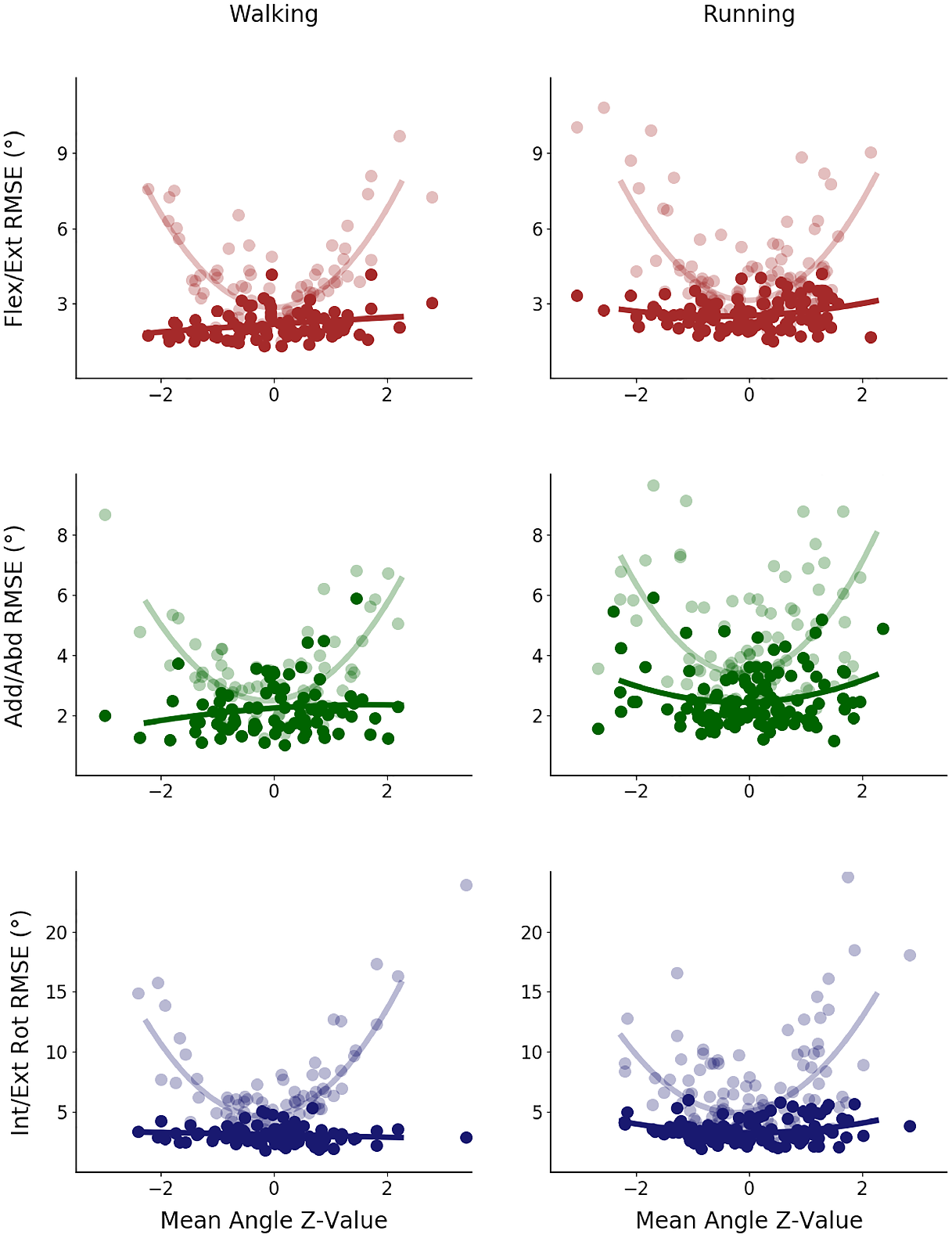
Effect of Passive Pseudo-Calibration. Prediction error for all three degrees of freedom was higher in subjects with overall kinematics that were further away from the mean, likely due to differences in limb alignment and sensor placement (light red, green, blue). Using the mean joint angles as a neutral pose, we passively calibrated joint angle predictions, which rendered the error relatively similar across subjects, despite their deviation from the mean (dark red, green, blue).

The accuracy of the models presented here is in part due to the large motion capture dataset that we used, which enabled the generation of a synthetic IMU database with hundreds of subjects. Large datasets are a requisite for training generalizable deep neural networks, but they are often not available in movement biomechanics. While the use of machine learning is growing, often deep neural networks are trained on limited data (Halilaj *et al*., 2018 [9]). Our experiments demonstrated that models trained on data from a larger number of subjects (i.e., hundreds) exhibit lower RMSEs compared to models trained on smaller data (i.e., tens), suggesting that they are more likely to generalize well to new, independent data. Learning curves (Fig. 4) are useful tools for determining how much data may be sufficient for a particular application, allowing one to weigh improvement in accuracy against efforts associated with acquiring more data. For studies designed with a machine learning approach in mind (instead of traditional hypothesis testing), they can replace power analyses in the planning phase.

Overall, this study introduced an integrated deep learning and optimization framework for the prediction of joint kinematics from inertial sensing data and presented a series of experiments that enable evaluation of the critical steps that contributed to the accuracy of these data-driven models. The presented techniques can also be extended to estimation of joint kinetics in the future (Johnson *et al*., 2020 [12]). While further validation of the models with true IMU data is a necessary next step, the promising results presented here, along with the publicly available tools, are encouraging steps toward addressing one of the most pressing technical challenges in modern-day biomechanics.

## 5. Acknowledgement

The authors would like to thank Allan Brett for his assistance with the data transfer and for patiently answering questions as the project progressed.

